# Diurnal Bull Ants Navigating by Moonlight: Polarised Light Homing in *Myrmecia tarsata*

**DOI:** 10.1101/2025.02.21.639436

**Authors:** Cody A Freas, Ken Cheng

## Abstract

We present evidence that the faint polarised moonlight pattern of the sky can be detected and used for navigation in the diurnal bull ant *Myrmecia tarsata*, despite this species lacking the highly refined low-light visual specialisations of the nocturnal bull ant, *Myrmecia midas*. The position of celestial bodies such as the sun and moon can provide navigating animals directional information, yet direct observation can often be occluded. Animals can estimate positional information of solar and lunar cues via their polarised light pattern, present across the sky. The sun’s polarisation pattern is widely used in animals and a similar, yet much fainter pattern is produced by the moon, yet it is unknown how widespread moonlight’s use is in navigating animals. Here, we demonstrate that the bull ant *Myrmecia tarsata*, which forages throughout the day, returning home at sunset, can use both solar and lunar polarised light patterns to navigate. We compare these findings to the closely related *M. midas* navigating under identical light conditions, as this nocturnal bull ant is known to rely on these polarised light patterns as part of their celestial compass. While *M. midas* and *M. tarsata* can clearly use both the solar and lunar polarised light patterns to navigate, *M. tarsata* showed degraded performance under polarised moonlight as a function of lunar phase, decreasing performance as illumination decreased. *M. midas* in contrast, exhibited impressive attendance to the overhead lunar polarisation pattern throughout the lunar month, illustrating the highly specialised low-light-detection adaptations of bull ants.

## Introduction

Polarised light describes light waves which oscillate in a directionally predictable fashion. When these waves oscillate along a single plane, they are defined as linearly polarised. Polarisation can arise naturally, with both sunlight and moonlight becoming polarised as a by-product of passing through the Earth’s upper atmosphere (Horváth & Varjú 2004; Horváth et al. 2014). The dispersion of this light forms a pattern across the sky, which is organised in concentric circles around the position of the celestial body it emanates form, creating an e-vector pattern. These circles radiate outward, extending up to 90° from their source, where this pattern both strengthens and becomes linearised (e-vector). Thus, when the sun or moon is near the horizon, they produce a linear polarisation pattern in the overhead sky, perpendicular to their position and roughly parallel to the north-south axis (Dacke et al. 1999, 2003; Reid et al. 2011; Zeil et al. 2014).

The polarisation pattern of the sun is present across the sky throughout the day and into twilight and is a useful compass cue for animal orientation and homing (Wehner and Müller 2006; Reid et al. 2011; Lebhardt and Ronacher 2013; Warrant and Dacke 2016; Freas et al. 2017a, 2019; Freas and Spetch 2023; Freas et al. 2024). Many animals navigate or hold a heading direction via the azimuthal position of the sun or moon, yet these can often be obscured by cloud cover or local vegetation (Jander 1957; Wehner and Lanfranconi 1981; Koltz and Reid 1993; Perez et al. 1997; Dacke et al. 2014; Warrant and Dacke 2016; Freas and Cheng 2022). Thus, the polarisation pattern provides a predictable proxy for where the sun and moon are positioned even when they are not directly visible (Horváth et al. 2014).

Insect navigators are well known to detect and attend to the polarisation pattern of the sky via polarisation-sensitive photoreceptors located in the dorsal rim area of the compound eye (Labhart and Meyer 1999; Homberg and Paech 2002; el Jundi et al. 2015) as well as their ocelli, which are also polarisation-sensitive (Fent and Wehner 1985; Fent 1986; Schwarz et al. 2011ab; Penmetcha et al. 2019). Much of what we know about polarised-light navigation focusses on diurnal animals utilising solar light polarisation, yet the moon also produces a similar yet much dimmer pattern radiating from its position, by reflecting sunlight off its surface (Gál et al. 2001). Multiple species of insects and other arthropods, including ants and bees, are known to track the moon’s observable position as part of their celestial compass (Jander 1957; Koltz and Reid 1993; Dacke et al. 2004; Ugolini et al. 2003), yet navigation via lunar polarised light remains understudied. Given that this polarisation pattern is much weaker than the sun’s, it has been shown to be observable by nocturnal animals which possess highly specialised eye physiologies for low light levels (Dacke et al. 2003, 2004, 2011; Smolka et al. 2016; Foster et al. 2019; Freas et al. 2024). Two species of nocturnal dung beetles (*Scarabaeus satyrus* and *S. zambesianus)* can attend to polarised moonlight for heading maintenance while they roll dung balls in a straight line away from a central dung pile (Dacke et al. 2003, 2004, 2011; Smolka et al. 2016; Foster et al. 2019). The large-eyed ants of the *Myrmecia* genus represent a group which posses highly attuned systems for navigating at low light (Narendra et al. 2016, 2017). Within this group, the nocturnal bull ant *Myrmecia midas* has recently been shown to use the polarised moonlight pattern to navigate, with the pattern used to update their path integrator overnight, just as diurnal ants use the sun’s pattern as part of their daytime celestial compass (Mote and Wehner 1980; Freas et al. 2024). Beyond these species, it has been hypothesised that this cue may be broadly used across nocturnal insect groups (Herzmann and Labhart 1989; Greiner et al. 2007; Rost and Honegger 1987).

Given that the moon creates such a dim version of the polarisation pattern formed around the sun, an initial prediction might be that highly specialised visual systems would be needed to reliably detect and attend to this pattern to orient and navigate to goals. Yet evidence suggests detection may be more widespread: the diurnal dung beetle *Scarabaeus* (*Kheper) lamarcki* also attends to moonlight polarisation patterns during their movement (Smolka et al. 2016), suggesting that moonlight may be a common cue for insect navigators at night regardless of their temporal activity niche. Though importantly, *S. lamarcki* does exhibit orientation performance decreases compared to their nocturnal counterparts, especially under crescent-moon-derived polarisation patterns (Smolka et al. 2016), hinting that lunar cues may not be useable to diurnal animals throughout the lunar cycle.

When exploring the possibility of widespread polarised-moonlight navigation use in insects, non-nocturnal species which possess some level of visual specialisation for dim light would be prime initial candidates for study. The diurnal/crepuscular bull ant *Myrmecia tarsata* exhibits many of these attributes. This species forages mainly during the day, yet its afternoon inbound foraging trip coincides with sunset and persists into the evening twilight (Greiner et al. 2007). Thus, foragers likely can navigate at least somewhat effectively in the declining light levels of twilight, relying on the solar polarisation pattern to inform their celestial compass, akin the diurnal/crepuscular desert harvester ant, *Veromessor pergandei* (Freas et al. 2019).

Physiologically, the eyes of *M. tarsata* represent somewhat of a mid-point between fully diurnal bull ants such as *Myrmecia croslandi* and nocturnal bull ants *M. pyriformis* and *M. midas*. Both facet lenses and photoreceptor diameters in *M. tarsata* foragers are larger than their counterparts in the diurnal *M. croslandi* yet are smaller than those in the nocturnal *M. pyriformis* (Greiner et al. 2007). *M. tarsata* and *M. pyriformis* also possess compound-eye modifications corresponding with low-light detection which are absent in diurnal bull ants (Narendra et al. 2016). The eyes of *M. tarsata* possess a variable proximal cone diameter which doubles in size in its dark-adapted state, though this effect is less pronounced compared to that of the nocturnal *Myrmecia pyriformis* (Narendra et al. 2016). Eye morphology comparisons between *M. tarsata* and the nocturnal *M. midas* have shown that *M. midas* had both larger ommatidia facet diameters and higher contrast sensitivity than *M. tarsata*, which should be important for *M. midas* navigating under low-light conditions. Importantly, low-light-detection specialisations can improve homing performance, both by enhancing the learning of terrestrial visual features of the environment as well as providing direct visual compass cues as part of these insects’ celestial compass. The presence of dim-light eye specialisations indicates that, while *Myrmecia tarsata* does not typically navigate under a lunar polarised sky, they may be able to detect this pattern when navigating at night. However, it also suggests that given their eyes are not as specialised as nocturnal bull ants (*M. pyriformis* and *M. midas)*, *M. tarsata* might struggle to match the performance of nocturnal bull ants when they attempt to navigate under the faint pattern of polarised moonlight.

In the current study, we compared the use of the overhead polarised-light pattern (both solar and lunar) in two sympatric bull ant species, the diurnal/crepuscular *M. tarsata* and the nocturnal *M. midas.* We characterised both species’ ability to orient under solar polarised-light cues, during evening or morning twilight (corresponding with each species’ natural inbound navigation periods), by rotating a linear polarising filter over them as they navigated to their nest (Figure 1). Additionally, we explored both species’ ability to detect and attend to polarised moonlight, during a full moon, when lunar cues are brightest, as well as across the lunar cycle, during quarter-moon and crescent-moon nights, when smaller portions of the moon’s surface reflect sunlight, producing a weaker lunar polarised light pattern. *M. tarsata* naturally returns to its nest during evening twilight, suggesting it likely naturally uses the solar polarisation pattern during this period to navigate. As this species is not typically active at night, it is unknown if it can detect and attend to lunar polarised light to navigate overnight. For a comparative perspective, we tested *M. midas* foragers during similar conditions as it is a well-studied nocturnal bull ant which we know uses both the overhead solar and lunar polarised-light patterns (Freas et al. 2017ab; Freas and Cheng 2018; Freas et al. 2024), even under crescent moons when the lunar polarisation pattern is at its faintest (Freas et al. 2024).

**Figure 1.**
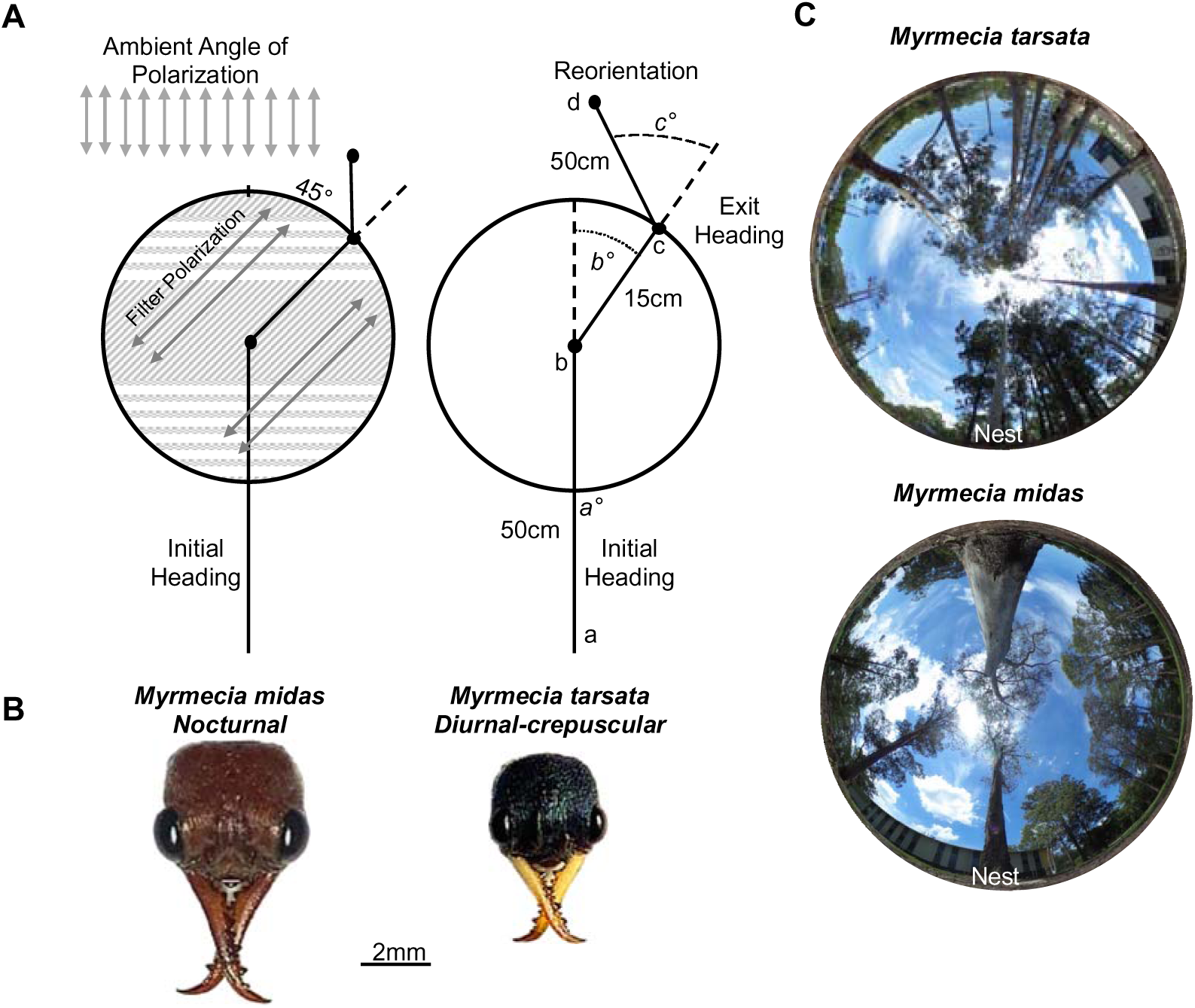
Diagram of the changes to the overhead polarisation pattern using the linear polarised filter during testing in *Myrmecia tarsata* and *M. midas*. (A)The ambient collar or lunar polarisation pattern, or e-vector, was altered by ±45°. Foragers initial heading directions, from release (a) to the site the filter was placed over each forager (b). Exit headings were calculated from the centre of the filter (b) to spot where the forager’s exit at the filter edge (c). Shifts under the filter (b°) were calculated taking the difference from the initial heading and exit heading. Reorientation were measured from the filter edge (c) to the forager’s heading 50cm post filter exit (d). Reorientation headings (c°) were calculated from the angular difference from the exit headings to reorientation heading direction. (B) Dorsal images of a forager’s head of each species. (C) Images of the sky and canopy cover at the two nests. Photos were taken at the on-route at the testing area for both nests.

## Methods

### Study site

Experiments were conducted from August 2024 to January 2025 on one *Myrmecia tarsata* nest and one *Myrmecia midas* nest on the Macquarie University campus in Sydney, Australia (33°46 11S, 151°06 40E). Both nests were located within eucalyptus tree stands with largely barren understories (Figure 1) and separated by 440m. *M. tarsata* is a largely diurnal forager, exhibiting two distinct bouts of foraging traffic, one in the morning with foragers returning before midday and one in the afternoon with foragers returning around sunset and into twilight (Greiner et al. 2007). In contrast*, M. midas* is fully nocturnal, with outbound foragers leaving the nest ∼20min after sunset, during evening twilight. These foragers travel along the ground to one of the surrounding trees and climb up into the canopy to feed overnight (Freas et al. 2017ab, 2018; Deeti et al. 2024). Inbound navigation in *M. midas* is more variable, with foragers descending their foraging tree and returning to the nest entrance both overnight and into morning twilight with activity ceasing by sunrise (Freas et al. 2017a). Both species are known to rely heavily on visual terrestrial cues while also accumulating a celestial-compass-derived path integrator for navigation (Freas et al. 2017ab, 2018; Isalm et al. 2020; Kamhi et al. 2020; Penmetcha 2023; Deeti et al. 2023).

### Linear Polariser Apparatus and Procedure

During testing, the overhead polarisation pattern was altered via placing a 30cm diameter linear polarisation filter (apparatus from Freas et al. 2024) above individual foragers as they navigated back to their nest. This filter blocks the ambient e-vector pattern of the night sky, replacing it with the e-vector orientation of the filter, ±45° the ambient pattern (Figure 1A). To allow foragers to travel beneath this filter, the linear polarising film was held in place by a rigid holding ring raised 10cm off the ground using four legs. Solar testing was conducted during evening or morning solar twilight coinciding with the natural inbound navigation of *M. tarsata* and *M. midas* (Figure 1B), respectively, while each lunar phase condition was tested overnight before solar twilight during a waxing moon (crescent, quarter and full). We obtained the sun rise and moon set positions as they reached the horizon based on Astronomical Almanac measurements (http://asa.usno.navy.mil)

Across all solar and lunar tests, held ants were released at their collection site and allowed to travel towards their nest for 50cm before the filter was placed overhead and all testing occurred within the first 3m from release (Figure 1C). Each ant was tested with both the ±45° e-vector rotation on a single inbound trip with the order randomised (either 1.) +45° then –45° or 2.) –45° then +45°). For the first test, we recorded four positions along the ant’s route, the site they were released, the ant’s position when the (±45°) filter was placed overhead, the ant’s position when they exited the filter ring, and their path ∼50cm after exiting the filter (reorientation heading). After reorientation, each ant was tested with the mirror e-vector-rotation filter orientation placed overhead. For this second test, we recorded their pre filter heading (50cm), the position when the filter was placed, the exit position and their position ∼50cm after exiting the filter (reorientation heading). The ant was followed during these periods using a redlight headlamp and stakes were used to mark the sites, taking care to not disturb the ant’s homing behaviour. These positions produced each forager’s initial heading under a natural e-vector pattern, their shift under an artificially rotated polarisation pattern, and their reorientation back to the natural pattern after exiting the filter (Figure 1A). After testing, each forager was recollected and marked with white acrylic paint (Tamiya™) to prevent retesting.

### Solar polarised light

For solar-polarised-light testing, foragers from both species were collected at the base of a foraging tree*. Myrmecia tarsata* foragers were collected from the ground ∼6m from their nest during their afternoon foraging period (4pm – 6pm), fed honey and held in a 10cm glass phial at their capture site until evening twilight testing (corresponding with the ending of their foraging activity). *Myrmecia midas* foragers were collected during evening twilight (the onset of their outbound foraging) at the base of their foraging tree (∼6m from the nest), just as they were about to climb onto the trunk. These foragers were fed honey and held in identical phials at the base of the foraging tree overnight and tested in the pre-dawn morning twilight, when the rising sun reached 15° below the horizon, and testing ceased at sunrise.

### Lunar Polarised light

Polarised-moonlight testing occurred during the lunar twilight, just after the setting moon crossed below the horizon until it reached 15° below the horizon. All testing occurred during waxing lunar phases, ensuring that the moon was present in the night sky from the end of solar twilight till testing, to reduce effects of overnight lunar-cue gaps (Freas et al. 2024). We tested both species under three waxing lunar phases, representing distinct amounts of lunar illumination: Full (>85%), Quarter (∼50%), and Crescent (∼25%). For all three lunar conditions, ants were collected identically to solar testing but released overnight, during moonset, when only the lunar polarisation pattern was present in the overhead sky. Given a true 100%-illumination full moon sets during morning solar twilight, when solar polarisation pattern cues would dominate, we tested during the two nights preceding the full moon when illumination was >85%, but when lunar twilight and morning solar twilight did not overlap.

### Statistical Analysis

Data were analysed with circular statistics with the statistics package Oriana (Batschelet, 1981; Fisher, 1993; Zar, 1999). Shift magnitudes under the filter were calculated by taking the difference between each forager’s initial heading and its heading as it exited the filter. Within-individual comparisons (initial heading vs. filter exit heading and filter exit heading vs. 50cm reorientation) were analysed using Moore’s Paired Tests. Across-species shift magnitudes were analysed using Watson–Williams F-tests between identical solar and lunar conditions (Solar, Full, Quarter, Crescent). To compare shift magnitudes between −45° and +45° both within a condition as well as any potential effect of experimental order, we mirrored the shifts (across the 0° to 180° plane) in each −45° condition for comparison with unaltered +45° shift data. We then compared these sets using Moore’s Paired Tests (Zar, 1999).

## Results

### Solar Polarised Light – Twilight Testing

In both *M. tarsata* (evening twilight testing) and *M. midas* (morning twilight testing), when the linear polarised filter was rotated clockwise (+45°) of the ambient pattern and placed above inbound foragers, filter exit orientations were significantly shifted from each forager’s initial heading (Table 1) predictably to the right of pre-filter headings (mean ± SE; *M. tarsata*: 34.7.0±9.5°; *M. midas*: 34.7±6.1°; Figure 2A,B). Foragers also predictably altered their headings when the filter was positioned counter-clockwise (–45°) the ambient e-vector. Exit headings in both species were to the left of initial pre-filter headings (*M. tarsata*: –37.0±6.9; *M. midas*: –36.7±9.9°; Figure 2A, B), and these shifts were significant (Table 1).

**Figure 2.**
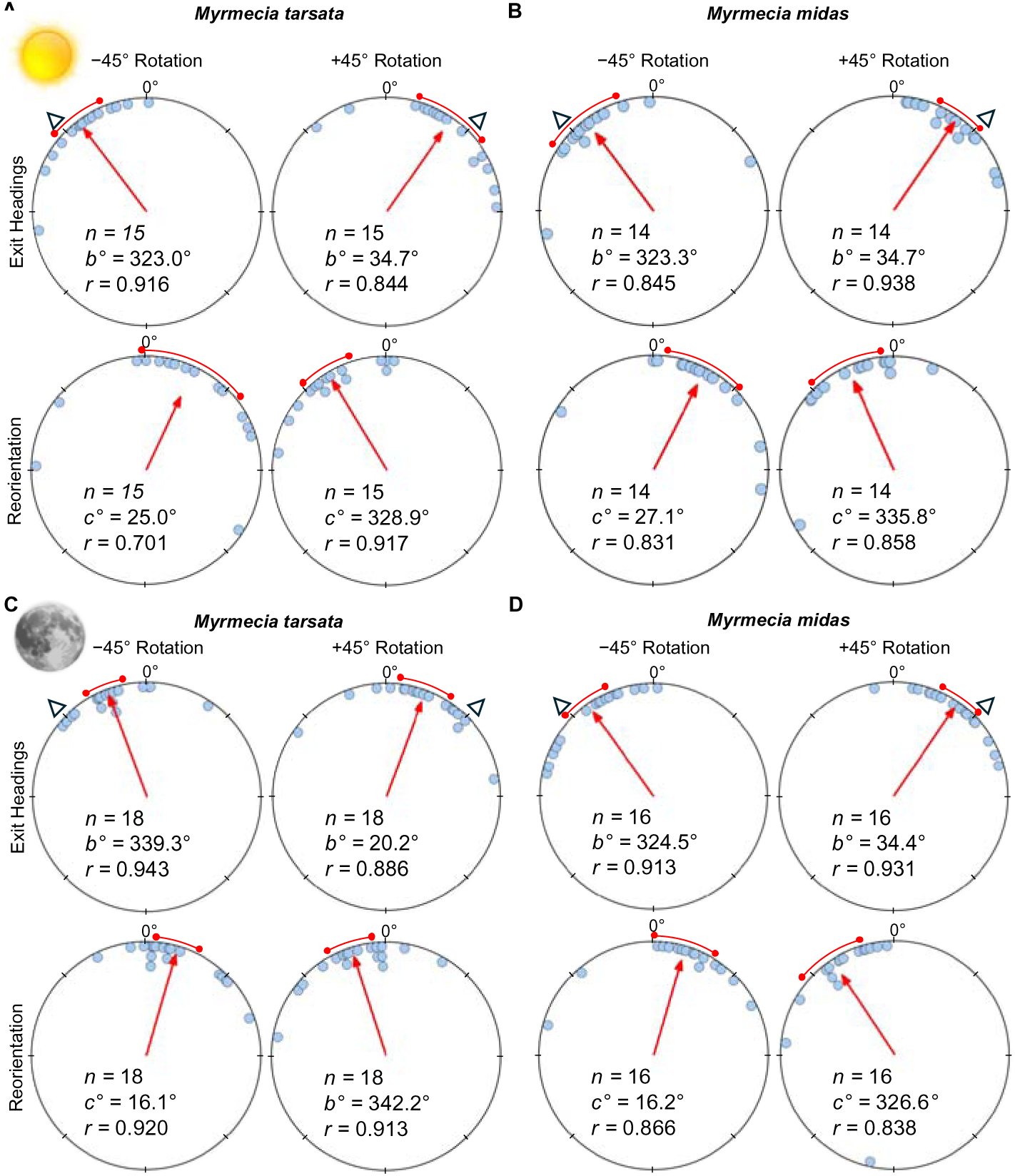
Circular distributions of headings during solar twilight and full moon conditions. In the solar condition, (A) *M. tarsata* and (B) *M. midas* were tested during their natural inbound twilight period, evening or morning twilight respectively. Lunar testing in (C) *M. tarsata* and (D) *M. midas* occurred during lunar twilight on the nights preceding the full moon (lunar illumination > 80%). Plots show the exit headings of foragers relative to their initial headings while the reorientation represents the change in headings 50cm after exiting the filter. Triangles indicate the ±45° e-vector change when forager are under the filter. Red arrows denote the length and direction of the mean vector of headings. *n*, number of individuals; *b*°, mean vector of shift; *c*°, mean vector of reorientation; *r*, length of the mean vector. Sun and moon images are public domain art accessed through wiki commons (https://commons.wikimedia.org/).

**Table 1.** Table showing the shift magnitudes and statistical comparisons for all tests.

|  | +45° Shift |  |  | −45° Shift |  |  | +45° Reorientation |  |  | −45° Reorientation |  |  |
| --- | --- | --- | --- | --- | --- | --- | --- | --- | --- | --- | --- | --- |
|  | mean<br>vector | Moore's Test |  | mean<br>vector | Moore's Test |  | mean<br>vector | Moore's Test |  | mean<br>vector | Moore's Test |  |
|  |  | R' | *p* |  | R' | *p* |  | R' | *p* |  | R' | *p* |
| Solar |  |  |  |  |  |  |  |  |  |  |  |  |
| *M. tarsata* | 34.7° | 1.496 | <0.001 | −37.0° | 2.004 | <0.001 | −31.1° | 1.600 | <0.001 | 25.0° | 1.343 | <0.005 |
| *M. midas* | 34.7° | 1.758 | <0.001 | −36.7° | 1.483 | <0.001 | −24.0° | 1.618 | <0.001 | 27.1° | 1.544 | <0.001 |
| Full Moon |  |  |  |  |  |  |  |  |  |  |  |  |
| *M. tarsata* | 20.2° | 1.431 | <0.005 | −20.7° | 1.089 | <0.05 | −17.8° | 1.485 | <0.005 | 16.1° | 1.290 | <0.01 |
| *M. midas* | 34.4° | 1.783 | <0.001 | −35.5° | 1.731 | <0.001 | −33.4° | 1.902 | <0.001 | 16.2° | 1.115 | <0.05 |
| Quarter Moon |  |  |  |  |  |  |  |  |  |  |  |  |
| *M. tarsata* | 14.3 | 1.298 | <0.01 | −18.1° | 1.59 | <0.001 | −19.2° | 1.244 | <0.025 | 22.8° | 1.29 | <0.01 |
| *M. midas* | 32.3 | 1.659 | <0.001 | −36.3° | 1.604 | <0.001 | −26.6° | 1.408 | <0.005 | 16.2° | 1.649 | <0.001 |
| Crescent Moon |  |  |  |  |  |  |  |  |  |  |  |  |
| *M. tarsata* | 3.8° | 0.276 | >0.500 | −4.0° | 0.639 | >0.100 | −3.0° | 0.207 | >0.500 | 17.0° | 0.643 | >0.100 |
| *M. midas* | 29.1° | 1.383 | <0.005 | −33.0° | 1.558 | <0.001 | 23.1° | 1.731 | <0.001 | 24.4° | 1.136 | <0.05 |

After exiting the (+45°) rotated filter, foragers of both species reoriented significantly (Table 1) to the left (mean ± SE; *M. tarsata*: –31.1±6.8°; *M. midas*: – 24.2±9.5°; Figure 2A, B). An similar significant shift (Table 1) to the right (mean ± SE *M. tarsata*: 25.0±13.8°; *M. midas*: 27.1±6.8°) was observed in both species after exiting the (–45°) rotated filter.

### Full Moon Lunar Polarised Light

Under a full moon (>80% illumination), when the polarisation pattern was altered by +45° or –45° of the ambient pattern, foragers of both species significantly (Table 1) altered their headings from pre-filter headings to the right (mean ± SE*; M. tarsata*: 20.2±6.6°; *M. midas*: 34.4±5.4°; Figure 2C,D) or left respectively (*M. tarsata*: – 20.7±5.5; *M. midas*: –35.5±6.1°; Figure 2C,D). In both species, after exiting the +45° or –45° rotated filter foragers reoriented significantly (Table 1) to the left (*M. tarsata*: –17.8±5.8°; *M. midas* –33.4±8.4°; Figure 2C, D) or right respectively (*M. tarsata*: 16.1±5.5°; *M. midas:* 16.2±7.7°).

### Quarter Moon

When tested under a waxing quarter moon and the overhead pattern was altered +45°, both species significantly (Table 1) altered their headings (mean ± SE, *M. tarsata*: 14.3±12.6°; *M. midas*: 32.3±5.7°). Opposite significant shifts (Table 1) to the left occurred when the overhead pattern was altered –45°, (*M. tarsata*: –18.1±5.1°; *M. midas*: –36.3±8.4°). After exiting the filter, forager reoriented significantly (Table 1) to the left (*M. tarsata*: –19.2±5.7°; *M. midas* –26.6±8.1°) or right, respectively (*M. tarsata*: 22.8±9.4°; *M. midas:* 16.2±7.7°).

### Crescent Moon

Under a waxing crescent moon, *M. tarsata* did not show significant shifts when the overhead pattern was altered ±45° (Table 1). In contrast and in line with previous work (Freas et al. 2024), *M. midas* still attended to the updated polarised light pattern under the filter under a crescent (∼25% illumination) moon. When the overhead pattern was altered +45° or –45°, M. midas foragers significantly (Table 1) altered their headings to the right or left respectively (+45°, mean ± SE, 29.1±12.1°; –45°, 33.0±9.3°).

### Across Species

Testing order within a homing trip between ±45° e-vector changes did not significantly alter shift magnitude within individuals in all conditions. Additionally, shift magnitudes between ±45° conditions (within individual) did not significantly differ in either *M. tarsata* or *M. midas* (*p>0.05*) across all conditions and were subsequently averaged, creating mean individual shifts for each solar and lunar condition. *Solar* mean individual shift magnitudes under an altered solar polarized light pattern (*M. tarsata*:35.7±7.2°; *M. midas*:35.5±6.9°) were almost identical and not significantly different between species (Watson-Williams F-Test; *F* _(1,27)_ = 0.001, p=0.975), suggesting similar weighting for solar polarised light cues in both species.

Lunar, *Full Moon:* Under lunar polarized light pattern, mean shifts magnitudes were significantly different between species (Watson-Williams F-Test; *F* _(1,32)_ = 6.154, p=0.019), with *M. midas* exhibiting larger individual shift magnitudes (*M. tarsata*:20.0±3.3°; *M. midas*:34.9±4.9°). Lunar, *Quarter Moon: M midas* also exhibited significantly larger (Watson-Williams F-Test; *F* _(1,22)_ =5.240, p=0.032) shift magnitudes under a quarter moon (*M. tarsata*:12.4±8.4°; *M. midas*:34.0±6.1°). Lunar, Crescent *Moon:* As would be expected, given *M. tarsata* did not exhibit significant shifts under a crescent moon, *M. midas* exhibited significantly larger shift magnitudes (Watson-Williams F-Test; *F* _(1,25)_ =13.94, p<0.001) during this lunar phase (*M. tarsata*:4.2±3.5°; *M. midas*:30.6±6.7°).

## Discussion

### Summary

These findings constitute the second instance of polarised-moonlight use for homing in insects. Both the nocturnal *M. midas* (Freas et al. 2024, replicated here) and the diurnal/crepuscular *M. tarsata* clearly attend to both solar (Freas et al. 2017) and lunar polarisation patterns when navigating home, evidenced by both species predictably altering their headings in line with rotations to the ambient e-vector while under the polarised filter. Our results indicate that solar and lunar polarised light can be detected and implemented into the celestial compass of diurnal/crepuscular insects, indicating that polarised-moonlight use may be widespread, even in groups which are not typically active overnight. Under an altered solar polarisation pattern during twilight, *M. tarsata* and *M. midas* were shown to alter their paths predicably under the filter, exhibiting almost the full 45° filter shifts, suggesting high levels of attendance to this solar cue for orientation during twilight. Under polarised moonlight, significant predictable heading shifts under the filter are still present in *M. tarsata* under full and quarter moons, but shift magnitudes declined along with smaller lunar phases, suggesting these foragers may have increased difficulty relying on this pattern as illumination decreases. Under a crescent moon, shifts became non-significant, indicating *M. tarsata* foragers could no longer attend to the polarised moonlight when it is at its weakest. In contrast, nocturnal *M. midas* showed only a marginal shift magnitude decline when tested under a lunar polarised light pattern, even under a *crescent* moon, suggesting this species can readily attend to this cue throughout the lunar month and corresponding with its highly specialised visual systems geared for low-light detection.

### Solar polarisation pattern

Nocturnal bull ants (*Myrmecia midas* and *Myrmecia pyriformis*) are known to use the solar polarisation pattern in the overhead sky to navigate during twilight periods (Reid et al. 2011; Freas et al. 2017ab), with *M. midas* being shown to rely on this pattern during both evening and morning twilight (outbound and inbound navigation, respectively). Current findings align with previous work in *M. midas* (Freas et al. 2017b) and reinforce that *M. midas* relies heavily on this solar pattern, updating their headings to changes in the overhead polarisation pattern almost fully to the altered filter e-vector. Foragers shifted their headings on average 35.7° (79% of the 45° solar e-vector rotation) while in Freas et al. (2017b) this shift was only marginally higher at 37.8° or 84% of the 45° solar e-vector rotation.

Solar polarised-light use in *M. tarsata* has not previously been studied, but their inbound navigation stretching into twilight suggested they were also likely to rely on this solar pattern to navigate, especially when the sun was not directly visible. This prediction was supported by work in another diurnal/crepuscular ant, the North American desert harvester ant (*Veromessor pergandei*), which attend to both the sun’s position and its polarisation pattern before sunset, weighting both equally. However, after sunset during evening twilight, these navigators switched to rely exclusively upon the solar polarisation pattern, almost fully shifting their headings under the filter 41.7° or 92.6% of the 45° solar e-vector rotation (Freas et al. 2019). As predicted, in the current study *M. tarsata* attended to the solar e-vector during evening twilight, updating their headings to the left or right when under the polarisation filter. Just as in *M. midas* and *V. pergandei*, *M. tarsata* shifts their headings almost fully, shifting 35.9° or 80% of the 45° solar e-vector rotation. These results reinforce that the solar polarisation pattern is readily detectable to both diurnal and nocturnal ant species and is likely a widespread compass cue when navigating during twilight.

A methodological explanation can be found for the propensity for foragers to underestimate the solar e-vector shift and subsequently only shifting their headings in response ‘almost fully’ (∼80-90%) across species and conditions, despite this being the brightest and most easily detectable celestial pattern (Reid et al. 2011; Freas et. 2017, 2019, 2024). As noted in previous work (Freas et al. 2024) and consistent with present testing, observations of the foragers when the filter is placed overhead suggest that heading shifts are not immediate. Instead, foragers continue in the original heading direction for a short distance before curving their path to the new direction. As the protocol measures the angle from the ant’s position at the filter centre when the filter is placed overhead rather than from the spot where the navigator altered its heading, this could cause measurements to slightly underestimate the shift.

### Lunar polarisation pattern

The typical description of nocturnal *Myrmecia* limiting their outbound foraging activity to evening twilight and then returning when light levels start to increase during morning twilight (Narendra et al. 2017) does not include the large amount of navigational activity that occurs overnight, when no solar cues are present to guide these ants. In both *Myrmecia pyriformis* and *Myrmecia midas*, foragers must conduct true nocturnal navigation as they return to the nest and leave to forage throughout the overnight period (Reid et al. 2011, 2013; Freas et al. 2017a). *M. tarsata,* despite foraging mostly diurnally, likely have some ability to detect faint light patterns such as the lunar polarisation pattern given its twilight navigation and attendance to solar polarisation after sunset. However, work on its eye sensitivity (Greiner et al. 2007; Narendra et al. 2016; Ogawa et al. 2019; Penmetcha et al. 2021) suggests it may struggle under a lunar pattern, especially when only small parts of the lunar plane are illuminated.

*M. midas* exhibited remarkably similar shifts under solar and lunar polarisation patterns, even under a crescent moon and in line with previous work (Freas et al. 2024). Under a full moon, foragers shifted almost the full 45° lunar e-vector manipulation (–45°: –35.5°/ +45°: 34.4°), representing 77.7% of the 45° filter rotation. These high shift magnitudes persisted throughout the lunar cycle with foragers shifting 76.2% of the 45° filter rotation under a quarter moon (–45°: –36.3°/ +45°: 32.3°) and 69.0% of the 45° filter rotation under a crescent moon (–45°: –33.0°/+45°: 29.1°). This represents only a ∼10% reduction in shift magnitude between the solar e-vector during twilight and the faintest e-vector of the crescent moon. Additionally, in all these conditions +45° or –45° was within the 95% confidence interval of heading shifts (Figure 2D; Figure 3BD) suggesting that *M. midas* foragers were fully attending to the updated polarisation pattern. Crescent-moon results in *M. midas* diverged slightly from previous work (Freas et al. 2024), with ants in the current study showing higher shift magnitudes under the same lunar phase (69.0% vs. 55.0%), but this difference was not statically significant.

**Figure 3.**
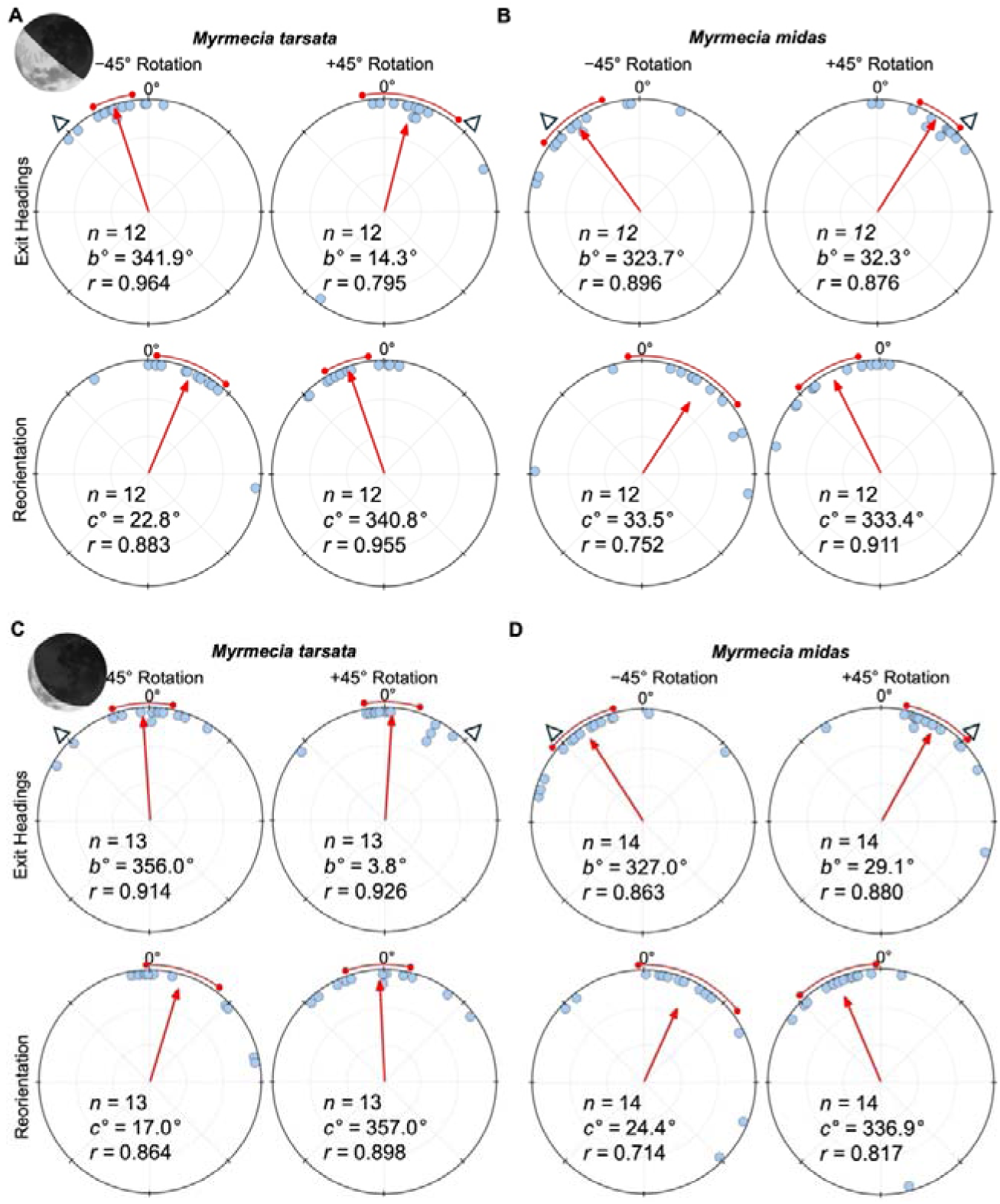
Circular distributions of headings during waxing quarter and crescent moon conditions. Under a quarter moon, (A) *M. tarsata* and (B) *M. midas* were tested during lunar twilight when the moon’s illumination was ∼50%. During crescent moon testing, in (C) *M. tarsata* and (D) *M. midas* occurred during lunar twilight on nights when the moon phase was a waxing crescent (lunar illumination ∼25%). Plots show the exit headings of foragers relative to their initial headings while the reorientation represents the change in headings 50cm after exiting the filter. Triangles indicate the ±45° e-vector change when forager are under the filter. Red arrows denote the length and direction of the mean vector of headings. *n*, number of individuals; *b*°, mean vector of shift; *c*°, mean vector of reorientation; *r*, length of the mean vector. Moon images are public domain art accessed through wiki commons (https://commons.wikimedia.org/).

In contrast, *M. tarsata’s* shift magnitudes decreased along with lunar phase, suggesting a reduced ability to detect the lunar e-vector pattern as light levels decrease. Under a full moon, *M. tarsata* foragers shifted under half of the 45° lunar e-vector manipulation (–45°: –20.7°/ +45°: 20.2°), representing 45.4% of the 45° filter rotation. Shift magnitude, while still significant, decreased further when tested under a quarter moon, with foragers only shifting 36% of the filter’s e-vector rotation (–45°: –20.7°/ +45°: 20.2°). Finally, *M. tarsata* foragers did not significantly shift under a crescent moon (–45°: –4.0°/ +45°: 3.8°), suggesting an inability to detect the faint lunar pattern produced by a crescent moon. Additionally, during crescent-moon testing, *M. tarsata* were extremely hesitant to move when under the filter, pausing as soon the filter was placed overhead for long periods, suggesting that when the surrounding landmarks are blocked via the filter, the crescent moon’s e-vector alone is not sufficient to guide homing. Statistically, across all three lunar conditions *M. tarsata* shifted significantly less than *M. midas*, suggesting these diurnal ants can detect and attend to the lunar polarisation pattern to home overnight but that the dim e-vector produced by the moon may be near this species’ light-detection threshold, resulting in smaller shifts when the pattern is altered via the filter, while the faint pattern present under crescent moons appears beyond their visual sensitivity.

Despite the performance decreases of polarised-moonlight detection in *M. tarsata* across lunar phases, these findings represent the second reported instance of this cue’s use in a diurnal animal (after the diurnal dung beetle *Scarabaeus* (*Kheper) lamarcki;* Smolka et al. 2016) as well as the second instance of its use as a tool in goal-directed navigation (after the nocturnal *M. midas* in Freas et al. 2024)

Interestingly, mirroring the differences and performance reductions in the current study, the diurnal *Scarabaeus* (*Kheper) lamarcki* exhibits similar reductions in performance under polarised-moonlight patterns compared to their closely related nocturnal species, *Scarabaeus satyrus* (Smolka et al. 2016). When the moon’s position was directly detectable by both species of dung beetle, they oriented equally as well while when presented only the lunar polarisation patten (as well as stars), diurnal dung beetles performed significantly worse than the nocturnal species, suggesting the dim-light adaptations in nocturnal insect eyes are necessary for precise orientation under faint light cues (Smolka et al. 2016).

## Conclusions

The diurnal/crepuscular bull ant *M. tarsata* can detect and attend to both the solar polarised light pattern, present at twilight, as well as the faint polarised moonlight pattern to home to the nest overnight, though not throughout the full lunar cycle. While *M. tarsata* exhibits similar performance to the nocturnal bull ant *M. midas* when tested under the solar polarisation patten, present when both are naturally active at twilight, *M. tarsata* shows reduced homing performance, represented by smaller shift magnitudes, under an altered lunar e-vector, compared to nocturnal bull ants. This performance degrades further with lunar phase-associated illumination decreases, suggesting that *M. tarsata* struggles to detect or attend to the faint moonlight pattern of quarter moon while the pattern produced under a crescent moon may be too faint to be usable for navigation. In contrast, the nocturnal *M. midas* exhibits only small decreases in performance under even the faintest of polarisation patterns produced under a crescent moon, suggesting the low-light-level visual adaptations possessed by nocturnal bull ants may be required for nocturnal navigation throughout the lunar cycle.

## Funding

This project was funded by a Macquarie University Research Fellowship (MQRF0001094), by Macquarie University, and by an ARC Discovery Grant (DP200102337).

## Land Acknowledgment

This work was conducted upon the Wallumattagal campus of Macquarie University. We acknowledge the traditional custodians of the land on which Macquarie University sits, the Wallumattagal clan of the Dharug Nation.

## Conflicts of interest

The authors declare no conflicts of interest associated with this work.

## Ethics

There are no state or federal governmental regulations guiding the research of invertebrates in Australia.

## Data Availability

All data, documentation and code will be made available online at osf.io. DOI:

## Notes

### Competing Interest Statement

The authors have declared no competing interest.

## References

1. Batschelet, E. (1981). Circular statistics in biology. Academic Press, 111 Fifth Ave., New York, NY 10003, 388.

2. Dacke, M., Byrne, M. J., Baird, E., Scholtz, C. H., & Warrant, E. J. (2011). How dim is dim? Precision of the celestial compass in moonlight and sunlight. Philosophical Transactions of the Royal Society B: Biological Sciences, 366(1565), 697–702.

3. Dacke, M., Byrne, M. J., Scholtz, C. H., & Warrant, E. J. (2004). Lunar orientation in a beetle. Proceedings of the Royal Society of London. Series B: Biological Sciences, 271(1537), 361–365.

4. Dacke, M., el Jundi, B., Smolka, J., Byrne, M., & Baird, E. (2014). The role of the sun in the celestial compass of dung beetles. Philosophical Transactions of the Royal Society B: Biological Sciences, 369(1636), 20130036.

5. Dacke, M., Nilsson, DE., Warrant, E. et al. (1999). Built-in polarizers form part of a compass organ in spiders. Nature, 401, 470–473. 10.1038/46773

6. Dacke, M., Nilsson, D. E., Scholtz, C. H., Byrne, M., & Warrant, E. J. (2003). Insect orientation to polarized moonlight. Nature, 424(6944), 33–33.

7. Deeti, S., Islam, M., Freas, C., Murray, T., & Cheng, K. (2023). Intricacies of running a route without success in night-active bull ants (*Myrmecia midas*). Journal of Experimental Psychology: Animal Learning and Cognition, 49(2), 111.

8. Deeti, S., Tjung, I., Freas, C., Murray, T., & Cheng, K. (2024). Heavy rainfall induced colony fission and nest relocation in nocturnal bull ants (*Myrmecia midas*). Biologia, 79(5), 1439–1450.

9. el Jundi, B., Warrant, E. J., Byrne, M. J., Khaldy, L., Baird, E., Smolka, J., & Dacke, M. (2015). Neural coding underlying the cue preference for celestial orientation. Proceedings of the National Academy of Sciences, 112, (36), 11395–11400.

10. Fent, K. (1986). Polarized skylight orientation in the desert ant Cataglyphis. Journal of Comparative Physiology A, 158, 145–150.

11. Fent, K., & Wehner, R. (1985). Ocelli: a celestial compass in the desert ant Cataglyphis. Science, 228(4696), 192–194.

12. Fisher, N.I. (1993). Statistical Analysis of Circular Data. Cambridge University Press, London. 10.1017/cbo9780511564345

13. Foster, J. J., Kirwan, J. D., El Jundi, B., Smolka, J., Khaldy, L., Baird, E., & Dacke, M. (2019). Orienting to polarized light at night–matching lunar skylight to performance in a nocturnal beetle. Journal of Experimental Biology, 222(2), jeb188532.

14. Freas, C. A., & Cheng, K. (2019). Panorama similarity and navigational knowledge in the nocturnal bull ant *Myrmecia midas*. Journal of Experimental Biology, 222(11), jeb193201.

15. Freas, C. A., & Cheng, K. (2022). The basis of navigation across species. Annual Review of Psychology, 73, 217–241.

16. Freas, C. A., & Spetch, M. L. (2023). Varieties of visual navigation in insects. Animal cognition, 26(1), 319–342.

17. Freas, C. A., Narendra, A., & Cheng, K. (2017a). Compass cues used by a nocturnal bull ant, *Myrmecia midas*. Journal of Experimental Biology, 220(9), 1578–1585.

18. Freas, C. A., Narenda, A., Murray, T., & Cheng, K. (2024). Polarised moonlight guides nocturnal bull ants home. eLife, 13, RP97615.

19. Freas, C. A., Narendra, A., Lemesle, C., & Cheng, K. (2017b). Polarized light use in the nocturnal bull ant, *Myrmecia midas*. Royal Society Open Science, 4(8), 170598.

20. Freas, C. A., Plowes, N. J., & Spetch, M. L. (2019). Not just going with the flow: foraging ants attend to polarised light even while on the pheromone trail. Journal of Comparative Physiology A, 205(5), 755–767.

21. Freas, C. A., Wystrach, A., Narendra, A., & Cheng, K. (2018). The view from the trees: nocturnal bull ants, Myrmecia midas, use the surrounding panorama while descending from trees. Frontiers in Psychology, 9, 16.

22. Gál, J., Horváth, G., Meyer-Rochow, V. B., & Wehner, R. (2001). Polarization patterns of the summer sky and its neutral points measured by full–sky imaging polarimetry in Finnish Lapland north of the Arctic Circle. Proceedings of the Royal Society of London A, 457(2010), 1385–1399.

23. Greiner, B., Cronin, T. W., Ribi, W. A., Wcislo, W. T., & Warrant, E. J. (2007a). Anatomical and physiological evidence for polarisation vision in the nocturnal bee *Megalopta genalis*. Journal of Comparative Physiology A, 193, 591–600.

24. Greiner, B., Narendra, A., Reid, S. F., Dacke, M., Ribi, W. A., & Zeil, J. (2007b). Eye structure correlates with distinct foraging-bout timing in primitive ants. Current Biology, 17(20), R879–R880.

25. Herzmann, D., & Labhart, T. (1989). Spectral sensitivity and absolute threshold of polarization vision in crickets: a behavioral study. Journal of Comparative Physiology A, 165, 315–319.

26. Homberg, U., & Paech, A. (2002). Ultrastructure and orientation of ommatidia in the dorsal rim area of the locust compound eye. Arthropod Structure & Development, 30(4), 271–280.

27. Horváth, G., & Varjú, D. (2004). Polarized light in animal vision: polarization patterns in nature. Springer Science & Business Media.

28. Horváth, G., Lerner, A., & Shashar, N. (2014). Polarized light and polarization vision in animal sciences (Vol. 2,). G. Horváth (Ed.). Berlin: Springer.

29. Islam, M., Freas, C. A., & Cheng, K. (2020). Effect of large visual changes on the navigation of the nocturnal bull ant, *Myrmecia midas*. Animal Cognition, 23, 1071–1080.

30. Jander, R. (1957) Die optische Richtungsorientierung der Roten Waldameise (Formica rufa L.) Zeitschrift tiir vergMehende Physiologie Bd, 40, 162–238.

31. Klotz, J. H., & Reid, B. L. (1993). Nocturnal orientation in the black carpenter ant Camponotus pennsylvanicus (DeGeer)(Hymenoptera: Formicidae). Insectes Sociaux, 40, 95–106.

32. Klotz, J. H., & Reid, B. L. (1993). Nocturnal orientation in the black carpenter ant Camponotus pennsylvanicus (DeGeer)(Hymenoptera: Formicidae). Insectes Sociaux, 40, 95–106.

33. Labhart, T., & Meyer, E. P. (1999). Detectors for polarized skylight in insects: a survey of ommatidial specializations in the dorsal rim area of the compound eye. Microscopy research and technique, 47(6), 368–379.

34. Lebhardt, F., & Ronacher, B. (2014). Interactions of the polarization and the sun compass in path integration of desert ants. Journal of Comparative Physiology A, 200, 711–720.

35. Mote, M. I., & Wehner, R. (1980). Functional characteristics of photoreceptors in the compound eye and ocellus of the desert ant, *Cataglyphis bicolor*. Journal of comparative physiology, 137, 63–71.

36. Narendra A, Greiner B, Ribi WA, Zeil J. (2016). Light and dark adaptation mechanisms in the compound eyes of Myrmecia ants that occupy discrete temporal niches. J Exp Biol 219:2435–42.

37. Narendra A, Kamhi JF & Ogawa Y. (2017). Moving in dim light: behavioural and visual adaptations in nocturnal ants. Integrative and Comparative Biology, 57, 1104–1116.

38. Ogawa, Y., Ryan, L. A., Palavalli-Nettimi, R., Seeger, O., Hart, N. S., & Narendra, A. (2019). Spatial resolving power and contrast sensitivity are adapted for ambient light conditions in Australian Myrmecia ants. Frontiers in Ecology and Evolution, 7, 18.

39. Penmetcha, B. (2023). Structure and physiology of the ant ocelli (Doctoral dissertation, Macquarie University).

40. Penmetcha, B., Ogawa, Y., Ribi, W. A. and Narendra, A. (2019). Ocellar structure of African and Australian desert ants. J. Comp. Physiol. A 205, 699–706.

41. Penmetcha, B., Ogawa, Y., Ryan, L. A., Hart, N. S., & Narendra, A. (2021). Ocellar spatial vision in Myrmecia ants. Journal of Experimental Biology, 224(20), jeb242948.

42. Perez, S., Taylor, O. & Jander, R. (1997). A sun compass in monarch butterflies. Nature, 387, 29. 10.1038/387029a0

43. Reid, S. F., Narendra, A., Hemmi, J. M., & Zeil, J. (2011). Polarised skylight and the landmark panorama provide night-active bull ants with compass information during route following. Journal of Experimental Biology, 214(3), 363–370.

44. Reid SF, Narendra A, Taylor RW & Zeil J. (2013). Foraging ecology of the night-active bull ant, Myrmecia pyriformis. Australian Journal of Zoology, 61, 170–177.

45. Rost, R., & Honegger, H. W. (1987). The timing of premating and mating behavior in a field population of the cricket *Gryllus campestris L*. Behavioral Ecology and Sociobiology, 21, 279–289.

46. Schwarz, S., Narendra, A. and Zeil, J. (2011). The properties of the visual system in the Australian desert ant Melophorus bagoti. Arthropod Struct. Dev. 40, 128–134.

47. Smolka, J., Baird, E., el Jundi, B., Reber, T., Byrne, M. J., & Dacke, M. (2016). Night sky orientation with diurnal and nocturnal eyes: dim-light adaptations are critical when the moon is out of sight. Animal behaviour, 111, 127–146.

48. Ugolini, A., Galanti, G., & Mercatelli, L. (2013). Do sandhoppers use the skylight polarization as a compass cue?. Animal Behaviour, 86(2), 427–434.

49. Warrant, E., & Dacke, M. (2016). Visual navigation in nocturnal insects. Physiology, 31(3), 182–192.

50. Wehner, R., & Lanfranconi, B. (1981). What do the ants know about the rotation of the sky?. Nature, 293(5835), 731–733.

51. Wehner, R., & Müller, M. (2006). The significance of direct sunlight and polarized skylight in the ant’s celestial system of navigation. Proceedings of the National Academy of Sciences, 103(33), 12575–12579.

52. Zar, J.H. (1999) Biostatistical Analysis. 4th Edition, Prentice Hall

53. Zeil J, Ribi WA, Narendra A. (2014). Polarization vision in ants, bees and wasps. In Polarized light and polarization visionin animal sciences, 2nd edn (ed.Gábor Horváth), pp. 41–60. Berlin, Germany:Springer Series in Vision Research.

